# Investigating high pathogenicity avian influenza virus incursions to remote islands: Detection of H5N1 on Gough Island in the South Atlantic Ocean

**DOI:** 10.1101/2025.09.06.674618

**Authors:** Antje Steinfurth, Joshua G. Lynton-Jenkins, Jaimie Cleeland, Benjamin C. Mollett, Holly A. Coombes, Andrea Moores, Robyn Neal, Ben Clifton, Marco Falchieri, Christopher W. Jones, Michelle M. Risi, Susannah Gold, Joe James, Peter G. Ryan, Jacob González-Solís, Ashley C. Banyard

**Author notes:** Corresponding author /.

## Abstract

Understanding the mechanisms underlying the emergence and spread of high pathogenicity avian influenza virus (HPAIV) is critical for tracking its global dissemination, particularly via migratory seabirds, given their role in transmission over long distances. Scavenging seabirds, such as skuas, may act as both reservoirs and vectors, and have been linked to multiple outbreaks since 2021. Here, we report the detection of HPAIV H5N1 clade 2.3.4.4b in three Tristan skua (*Stercorarius antarcticus hamiltoni*) carcasses on Gough Island in the central South Atlantic Ocean. To investigate potential incursion routes, we combined genomic analyses with year-round tracking data from global location sensors. Although migratory movement patterns suggested southern Africa as the most obvious pathway, the strain detected on Gough Island was more closely related to that identified in South Georgia, indicating that infection may have occurred during the pre-laying exodus, when skuas disperse into frontal waters south of the island. No further cases have been confirmed for Gough, but more systematic monitoring is needed to understand the dynamics of virus infection. The detection of HPAIV H5N1 in skuas on Gough Island highlights the importance of continued vigilance, proactive and geographically inclusive surveillance strategies, and biosecurity measures globally, alongside efforts to reduce other pressures on globally important seabird populations to help strengthen their resilience.

## Introduction

The ongoing expansion of high pathogenicity avian influenza virus (HPAIV) poses a major threat to wildlife, livestock, and, through its zoonotic potential, human health. Since first emerging in poultry in China in 1996, the A/goose/Guangdong/1/96 (GsGd)-lineage of HPAIV H5Nx has evolved to spread efficiently among a broad range of bird species globally, causing mass mortality events in unprecedented numbers [1–6]. Following its dissemination across Europe by migratory species, H5N1 reached North America via the East Atlantic flyway and, by October 2022, had expanded rapidly throughout South America, with devastating impacts on both seabird and pinniped populations [7,8]. In the austral spring of 2023, HPAIV reached the sub-Antarctic islands and Antarctica, first detected in brown skuas (*Stercorarius antarcticus*) on Bird Island, South Georgia [9]. Soon after, cases were reported in southern fulmars (*Fulmarus glacialoides*) and black-browed albatrosses (*Thalassarche melanophris*) in the Falkland Islands [9], and in multiple species across the Antarctic Peninsula [10]. Genetic analyses indicate that the detections in these regions were linked to strains originating in South America, likely introduced via long-distance movements of scavenging seabirds such as skuas, gulls, and giant petrels [9, 11]. In the following breeding season 2023/2024, the virus had spread further east across the sub-Antarctic and into the southern Indian Ocean, causing significant die-offs in wandering albatross (*Diomedea exulans*) chicks on Marion Island [12], before it was reported for the Crozet and Kerguelen Archipelagos, where mass mortality events predominantly affected southern elephant seals (*Mirounga leonina*) [11]. Genetic evidence suggests that the Crozet and Kerguelen outbreaks resulted from independent introductions linked to South Georgia rather than from the nearer southern African coast [11].

Gough Island in the central South Atlantic is situated approximately 3,600 km northeast of South Georgia and 2,450 km northwest of Marion Island and considered one of the most important seabird breeding sites globally [13]. Despite its small size (∼65 km²), the island supports an estimated eight million birds of at least 24 species, many of them endemic or near-endemic to the island or island group of Tristan da Cunha, and several of global conservation concern, including the Critically Endangered Tristan albatross (*Diomedea dabbenena*), MacGillivray’s prion (*Pachyptila macgillivrayi*), and Gough finch (*Rowettia goughensis*) [13–16]. Consequently, the potential for the arrival of HPAIV into this sensitive environment has remained a significant conservation concern. A risk assessment in 2022 considered the likelihood of the emergence of HPAIV on Gough Island to be ‘low’ [17]. However, the spread of the HPAIV epizootic into the southern hemisphere and its continued expansion across the sub-Antarctic region have substantially increased the risk of incursion.

Recurrent HPAIV outbreaks have also been reported in coastal areas of southern Africa, following viral spread from Europe via the Afro-Eurasian flyway [18–20]. Therefore, Gough Island’s unique geographical position in the South Atlantic along major migratory flyways and marine corridors, linking the Atlantic and Southern Oceans, renders it vulnerable to HPAIV introduction from multiple directions [21].

Antibodies against AIVs in Gough’s breeding seabird population were first detected in 2009, when a serosurvey on Gough Island revealed exposure in brown skuas (*S. a. hamiltoni;* hereafter referred to as Tristan skua) and northern rockhopper penguins (*Eudyptes moseleyi*), demonstrating that influenza A viruses have previously reached the island and circulated among these species [22, 23]. The Tristan skua is endemic to the island group of Tristan da Cunha, with most of its population breeding on Gough [24]. Given their scavenging behaviour, skuas represent both key reservoirs and vectors for disease transmission, making the study of their movements critical for identifying introduction pathways [23, 25].

Here, we describe the detection of HPAIV H5N1 clade 2.3.4.4b in three Tristan skuas from Gough Island in September 2024 and investigate potential incursion routes by using a combination of genetic analyses of the virus and year-round tracking data from global location sensors.

## Materials & Methods

### Study site

Gough Island (40°19′12″S 09°56′24″W) is part of the British Overseas Territory of St Helena, Ascension, and Tristan da Cunha in the central South Atlantic Ocean (Fig.1). Gough Island lies approximately 380 km south-southeast from the Tristan da Cunha archipelago, which consists of three main islands: Tristan da Cunha (96 km²), Inaccessible (14 km²), and Nightingale (4 km²), along with its satellite islets Middle (or Alex) and Stoltenhoff (both ∼0.1 km²), all within 40 km of each other (Fig. 1). Gough is uninhabited except for a small year-round team of 7-9 people operating the research station. The station is resupplied once a year in September/October by the South African research vessel *SA Agulhas II*. A field team of 2-3 biologists is on the island year-round as part of the island’s long-term term monitoring and science programme, trained to provide surveillance for unusual avian mortality events.

**Figure 1.**
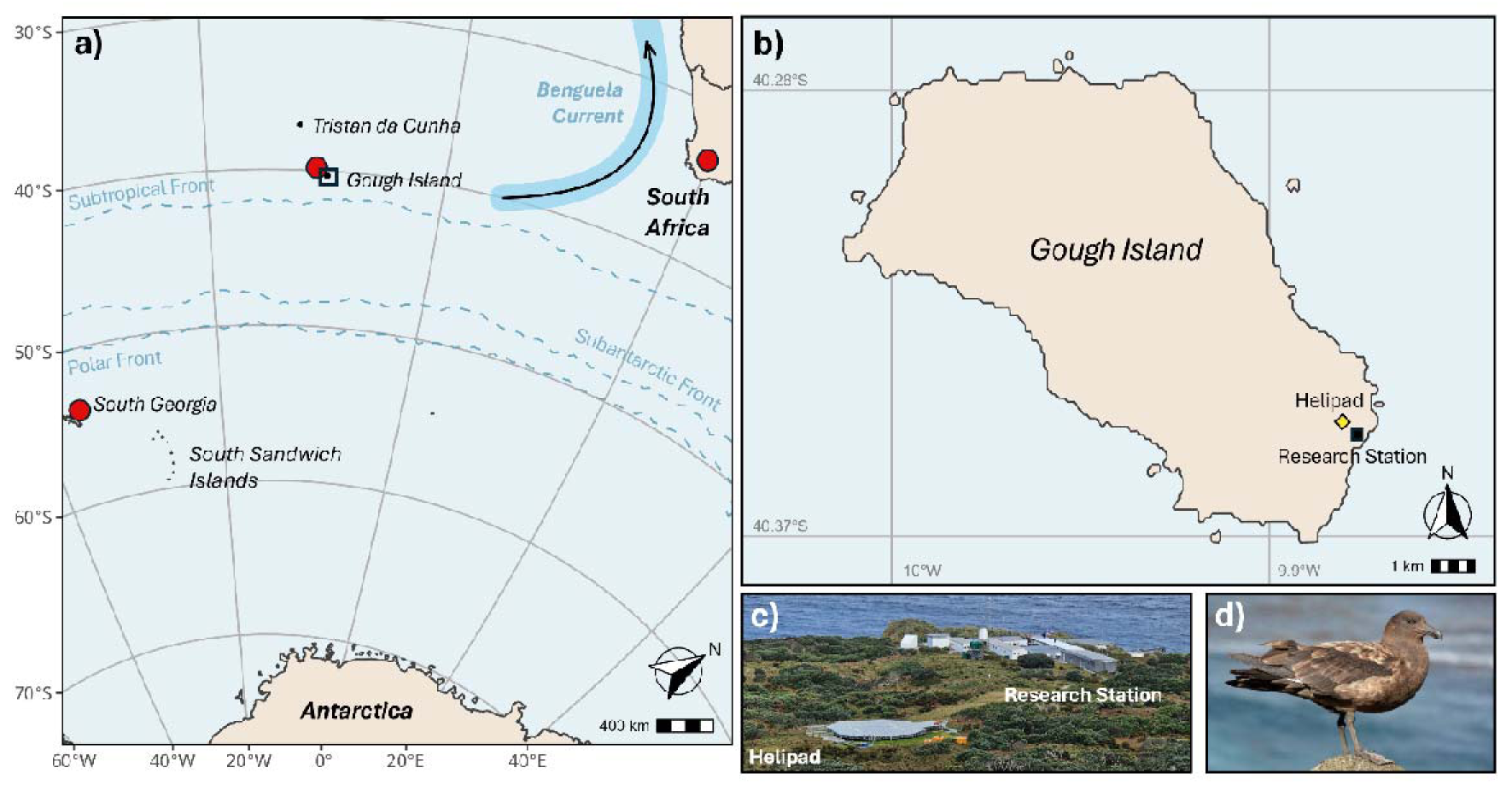
a) Map indicating the location for Gough Island in the context of locations with ongoing HPAIV H5N1 outbreaks reported to WAHIS (red dots), b) Gough Island with the research station located in the southeast of the island (black square) and the island helipad’s (yellow diamond), c) view of research station and helipad, d) Tristan skua (*Stercorarius antarcticus hamiltoni*).

### Tracking data

Migrate Technology Intigeo C330 geolocation tags (17 x 19 x 8 mm, 3.3 g) were deployed on 10 incubating adult Tristan skuas near the Gough Island research station in October-November 2017. The tags were attached to standard stainless steel SAFRING rings. Eight tags were retrieved during the following breeding season; the two other devices were not recovered.

Light-level data were processed in R version 4.4.1 using the *SGAT* package (v0.1.3) [26], implementing a Bayesian state-space model to estimate probable locations based on the timing of twilight transitions, while accounting for uncertainties in light detection and movement behaviour. Raw light data were pre-processed using the *BAStag* package (v0.1.3) [27] to identify twilight events (sunrise and sunset) based on light intensity thresholds. These twilight times were then manually reviewed and edited to correct for anomalies caused by shading, cloud cover, or behavioural factors e.g., incubation. The resultant data were used to infer two positions per day, corresponding to daily twilight transitions.

To quantify space use, we calculated kernel utilisation distributions (KUDs) using the *adehabitatHR* package (v0.4.22) [28]. *SGAT*-derived location estimates were projected onto an equal-area coordinate reference system to allow for unbiased spatial analysis. For each bird, 50% and 95% KUDs were calculated to represent core use areas and home ranges, respectively, across the pre-laying, breeding, and non-breeding periods. Location data during incubation in 2017 and 2018 were pooled for KUD analysis. To evaluate population-level space use, individual KUDs were averaged to assess spatial habitat use. Migration pathways between breeding and non-breeding areas were excluded from the KUD analysis and instead visualised using raw location data. For each individual, the onset of north-east migration away from the breeding colony and that of south-west return migration was defined as the point at which the individual’s distance from the colony increased or decreased continuously, without subsequent decreases/increases. The end point of each migration phase was defined as the point at which the distance from the colony stabilised, indicating arrival at a new residency area. All spatial operations were conducted using the *sp*, *sf*, and *raster* packages in R.

While we acknowledge that the GLS data used in this study were collected up to six years prior to the HPAI detection, we expect movement patterns of this population to have remained largely consistent. Long-term (multiyear) fidelity to specific wintering sites is common in migratory seabirds, particularly when these sites are associated with predictable environments such as upwelling systems, where repeated use can increase familiarity with local resources and prey dynamics [29–31]. Although climate change may gradually alter the predictability of marine environments [32,33], the persistence of the wintering site used by Tristan skuas across multiple breeding seasons (2009/10 (5 tracks), 2010/11 (9), 2011/12 (6), and 2017/18 (8)) spanning nearly a decade [34, this study], suggests that this fidelity continues to confer ecological benefits and thus is likely representative of the wider Gough Island skua population today.

### Sample collection

Swab samples from the brain, trachea, and cloaca of dead birds were collected, frozen at −20°C within one hour of collection, and shipped to the World Organisation for Animal Health/Food and Agriculture Organisation (WOAH/FAO) International Reference Laboratory for Avian Influenza at the Animal and Plant Health Agency (APHA) in Weybridge, UK, for diagnostic evaluation. Surveillance for suspicious wildlife mortality continued with special attention given to any spatial or temporal clustering of cases or unusual behaviours in wildlife.

### Molecular methods

Total nucleic acid was extracted from all swab samples as described previously [35] and viral RNA was screened using four real-time reverse transcription polymerase chain reaction (rRT-PCR) assays including: i) a Matrix (M)-gene assay for generic influenza A virus detection [36], ii) a HPAIV H5 clade 2.3.4.4b specific assay for HA subtype and pathotyping [35], iii) an N1-subtype specific rRT-PCR to confirm the neuraminidase type [37], and iv) an avian paramyxovirus type-1 (APMV-1) large polymerase (L)-gene assay [38]. All rRT-PCRs were undertaken on the AriaMx qPCR System (Agilent, United Kingdom). Material from positive brain tissue swabs was used for virus isolation in 10-day-old specific pathogen-free (SPF) embryonated fowls’ eggs according to internationally recognised methods (Codes and Manuals - WOAH - World Organisation for Animal Health).

### Viral Sequencing

Whole-genome sequencing (WGS) was undertaken on brain tissue swabs. Extracted vRNA was converted to double-stranded cDNA and amplified using a one-step RT-PCR using SuperScript III One-Step RT-PCR kit (Thermo Fisher Scientific) as previously described [39]. PCR products were purified with Agencourt AMPure XP beads (Beckman Coultrer) prior to sequencing library preparation using the Native Barcoding Kit (Oxford Nanopore Technologies) and sequenced using a GridION Mk1 (Oxford Nanopore Technologies) according to manufacturer’s instructions. Assembly of the influenza A viral genomes was performed using a custom in-house pipeline as previously described [9].

### Phylogenetic Analysis

To identify the genetic relatedness of Gough Island viruses to other reported strains, HPAIV H5N1 clade 2.3.4.4b sequences from Europe, Africa, Antarctica and South America available in the EpiFlu database between 1^st^ January 2020 and 28^th^ July 2025 were collated. To remove over-represented groups, the sequences were subset to cover 0.5% sequence divergence using PARNAS [40]. The remaining dataset was separated by segment and aligned using Mafft v7.525 [41] and manually trimmed to the open reading frame using Aliview v.2021 [42]. The trimmed alignments were used to infer maximum-likelihood (ML) phylogenetic trees using IQ-Tree v2.4.0 [43] with model finder, 1000 ultrafast bootstraps and SH-like approximate likelihood ratio test. Clade classification of Gough Island sequences was confirmed with GenoFLU-multi https://github.com/moncla-lab/GenoFLU-multi.

To produce a time scale phylogeny, all H5 HA sequences from South America and Antarctica were used alongside a subset of North American sequences to improve temporal signal. The dataset was aligned, trimmed, and an ML phylogenetic tree was inferred using the same approaches as described above. The HA ML phylogeny was then used to infer a time scaled phylogeny using TreeTime v0.11.4 [44]. Discrete trait analysis of transitions between locations was carried out using the mugration inference model in TreeTime with default settings.

## Results

The first suspected HPAIV-related death of an adult Tristan skua was observed on Gough Island on 12 September 2024 at a club of 50-200 non-breeding skuas at the helipad near the research station. The following day, another skua at the same location exhibited clinical signs consistent with HPAIV, including drooping head and wings, lethargy, and an inability to walk or fly, and was found dead later the same day [45]. Two additional carcasses were discovered on 15 September. Swab samples were collected from three of the four carcasses on 20 September 2025, and the carcasses were buried afterwards. The fourth carcass had disappeared by the time of sampling and was presumed to have been scavenged by other skuas.

### Diagnostic evaluation of samples

All three birds sampled on Gough Island tested positive for HPAIV H5N1 in oropharyngeal and brain swabs but tested negative in cloacal swabs (Table 1). Brain samples indicated a high viral load with HPH5 rRT-PCR values ranging from 19.5 - 23.8 Cts. Virus isolation was attempted on two of the three brain samples and both yielded hemagglutinating virus. Whole genome sequences (WGS) obtained from two samples confirmed the viruses belonged to HA clade 2.3.4.4b, genotype B3.2, as previously detected in the Antarctic region (Fig. 2a, Supplementary Fig. S1) [9]. The two WGS shared >99% identity across all genes. Phylogenetic analysis shows that these viruses are part of a monophyletic clade within the Antarctic Peninsula and sub-Antarctic region, including ancestral viruses from South Georgia and their descendants in Crozet and Kerguelen (Fig. 2) which is consistent with an expansion of the introductory pathway described by Banyard et al. [9], rather than a new incursion from Africa. Analysis of the asymmetric transition rate matrix reveals notable patterns of viral movement among key sub-Antarctic and South Atlantic islands (Fig. 2b). South Georgia appears to be a significant ancestral source, exhibiting outward transitions to Crozet, Kerguelen, Gough Island and the Falkland Islands. Gough Island demonstrates moderate connectivity across the region, particularly with Crozet and Kerguelen. However, transition rates from Gough Island to South Georgia and back suggest less frequent bidirectional exchange. Overall, the data supports a model in which South Georgia serves as a primary source, seeding transmission across the sub-Antarctic and South Atlantic islands (Fig. 2b).

**Figure 2.**
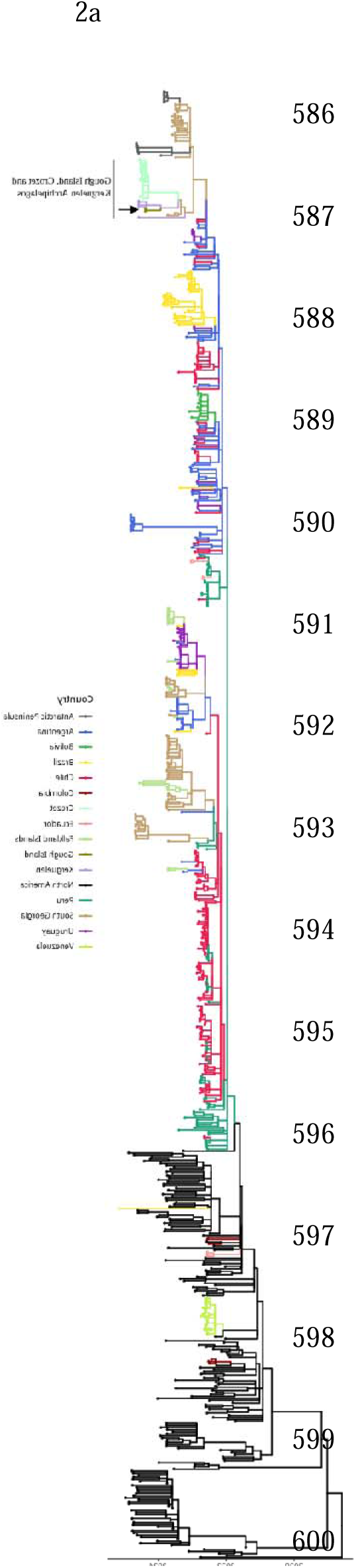
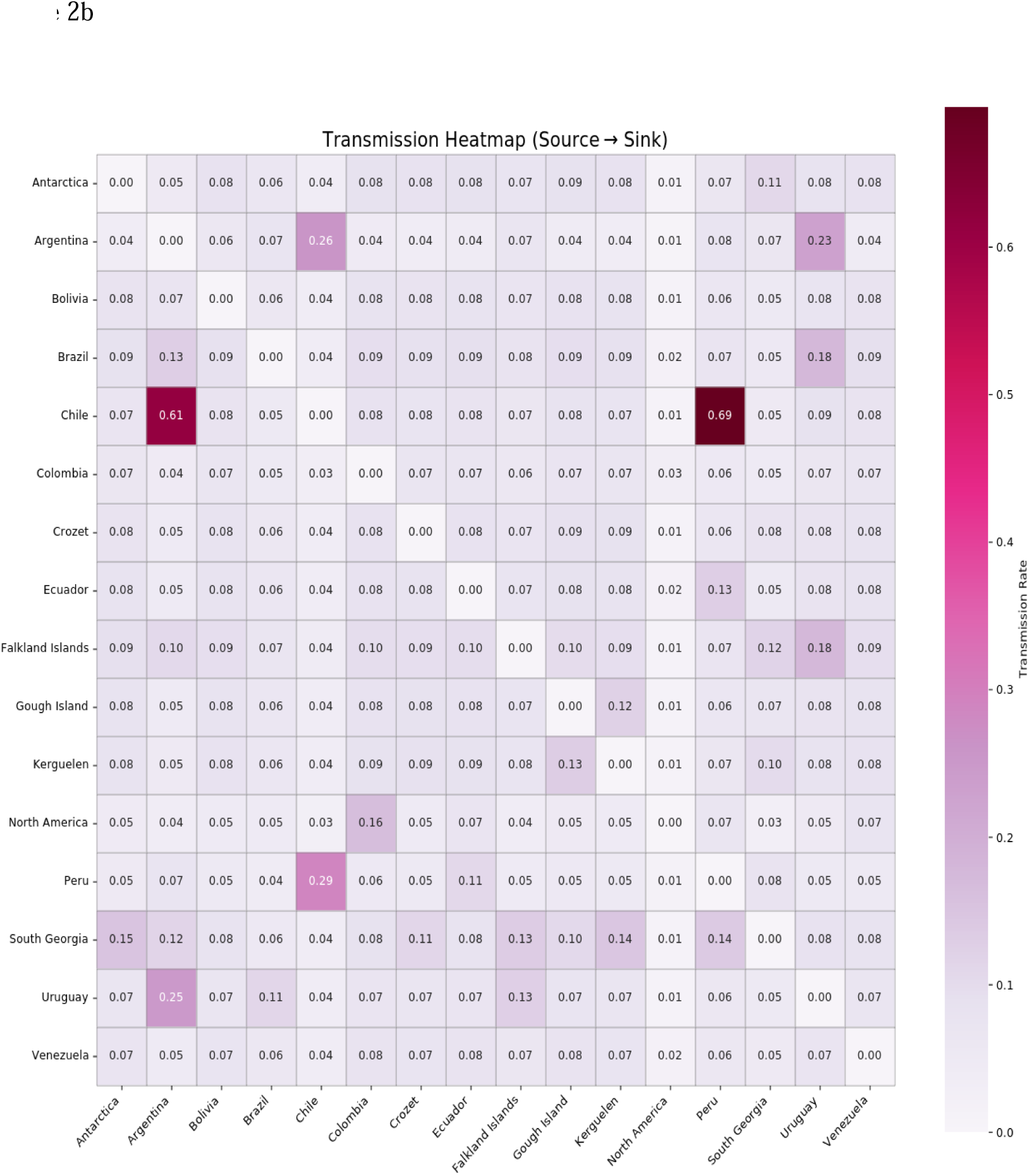
a) Maximum-likelihood tree of HA-gene including the two Gough Island sequences identified from this study (indicated by an arrow) and 944 HA-gene sequences from Antarctica, South and North America previously identified as 2.3.4.4b. Sequences from South America are coloured according to country, all sequences from North America in black, b) a source-sink heat map of transmission rate estimates between geographic regions.

**Table 1:** RT-PCR results, virus isolation and viral whole genome sequencing on samples collected from three Tristan skuas (*Stercorarius antarcticus hamiltoni*) on Gough Island on the 20th September 2024.

| Bird number | Sample Type<br>(Swab) | M Gene<br>RT-PCR | H5HP<br>RT-PCR | N1<br>RT-PCR | Interpretation | Strain | GISAID<br>Accession<br>number | Isolate |
| --- | --- | --- | --- | --- | --- | --- | --- | --- |
| 1 | Tracheal | Positive | Positive | Positive | HPAIV H5N1 | A/Tristan_Skua/Gough_Island/047354/2024_\|H5N1\|_2024<br>-09-20 | EPI_ISL_201<br>51596 | Yes |
|  | Cloacal | Negative | Negative | Negative |  |  |  |  |
|  | Brain | Positive | Positive | Positive |  |  |  |  |
| 2 | Tracheal | Positive | Positive | Positive | HPAIV H5N1 | A/Tristan_Skua/Gough_Island/047355/2024_\|H5N1\|_2024<br>-09-20 | EPI_ISL_201<br>51597 | NA |
|  | Cloacal | Negative | Negative | Negative |  |  |  |  |
|  | Brain | Positive | Positive | Positive |  |  |  |  |
| 3 | Tracheal | Positive | Positive | Positive | HPAIV H5N1 | A/Tristan_Skua/Gough_Island/<br>050656/2024_\|H5N1\|_2024-09-20_EP1 | NA | Yes |
|  | Cloacal | Negative | Negative | Negative |  |  |  |  |
|  | Brain | Positive | Positive | Positive |  |  |  |  |

### Ringing and tracking data

Tracking durations ranged from 343 to 353 days (mean ± SD: 345 ± 3.3 days), yielding 687 ± 7.8 locations per individual (Fig. 3 and Fig. 4). Kernel utilisation distributions revealed clear seasonal shifts in space use by Tristan skuas (Fig. 4a). During the pre-laying exodus, birds concentrated their foraging activity in waters surrounding the Subantarctic front, with a mean 95% utilisation distribution (home-range) of 977,900 km² and a corresponding 50% core area of 151,100 km². During the breeding period, space use became more restricted around Gough Island, with a reduced mean 95% UD of 567,000 km² and a 50% core area of 128,900 km², consistent with central-place foraging constraints. In contrast, the non-breeding period was characterised by a marked expansion and displacement of activity areas to the South African and Namibian continental shelves, where skuas occupied a mean 95% UD of 1 711,300 km² and a 50% core area of 394,400 km². Across these periods, core areas consistently represented a smaller, high-use subset of the broader home-range, while migration corridors were excluded from KUD estimation and are presented separately as movement tracks. Following the breeding season, individuals departed the foraging grounds around Gough Island between 12 January and 12 February 2018, *en route* to non-breeding foraging grounds off the coasts of South Africa and Namibia. Arrival at these non-breeding areas occurred between 20 January and 25 February 2018, with the north-eastward migration taking 11.9 ± 4.9 days (range 4–17 days). Individuals remained within the non-breeding wintering grounds for 141 ± 19.5 days (range 109–165 days). Departures from these areas occurred between 4 June and 15 July 2018, with arrival back at the pre-laying foraging area near Gough Island occurring between 13 June and 1 August 2018. The south-westward return migration took 13.8 ± 6.1 days (range 7–23 days).

**Figure 3.**
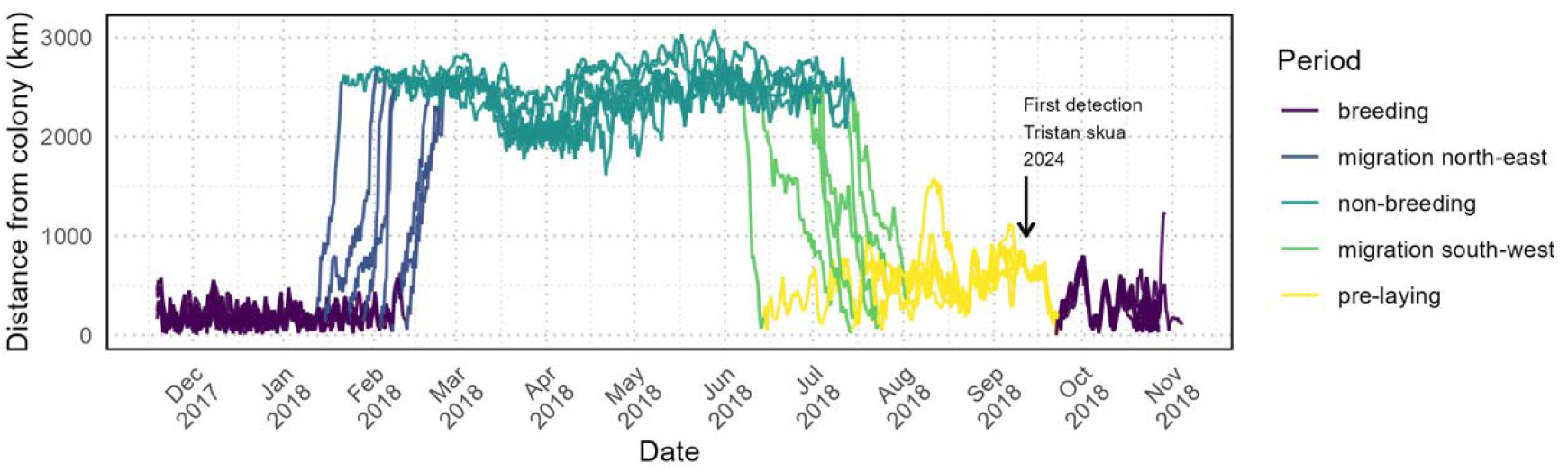
Time series showing the distance from the colony of GLS-tracked Tristan skuas (8) during the 2017/18 and 2018/19 breeding seasons, across key stages of the breeding cycle.

**Figure 4.**
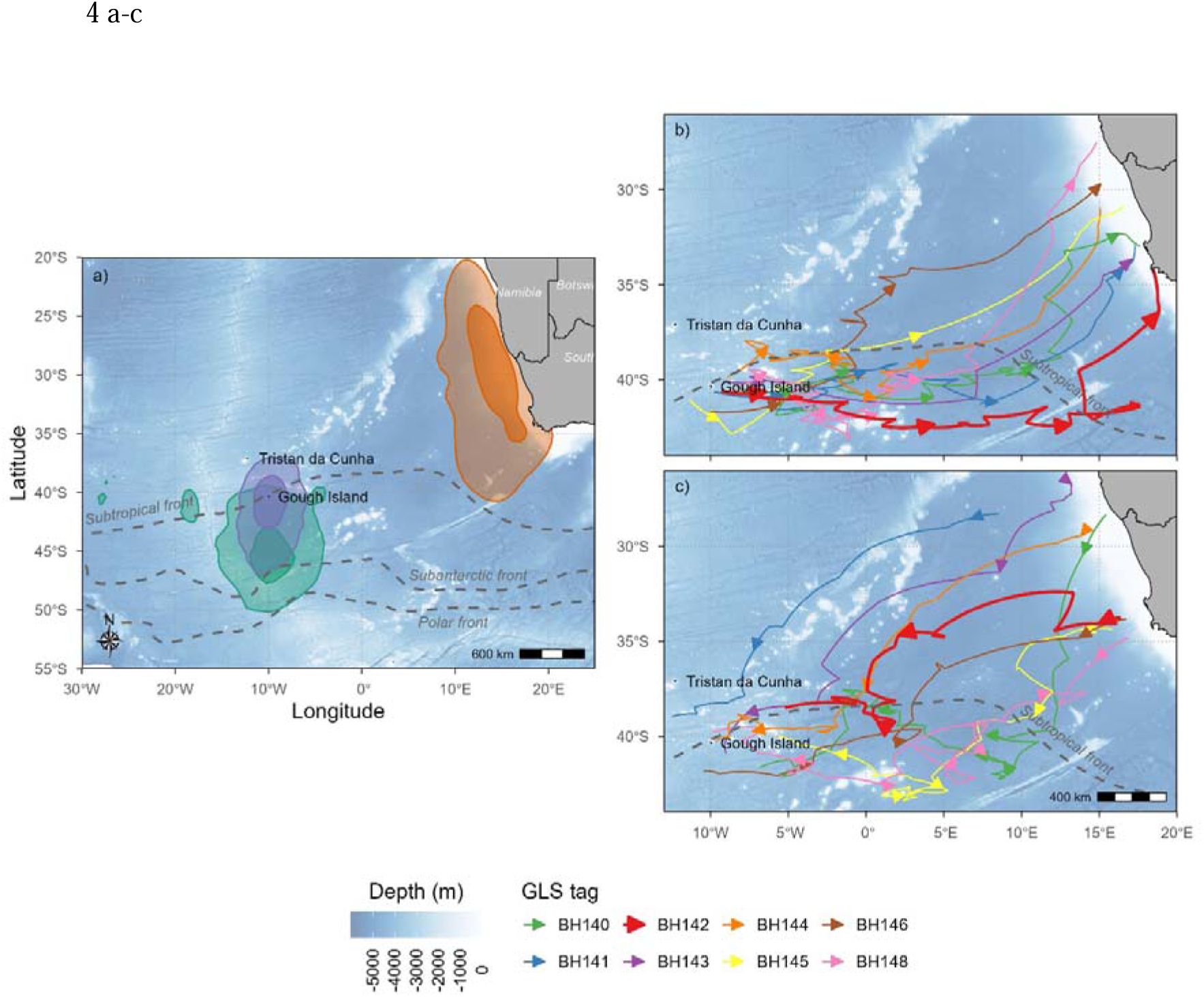
Kernel utilisation distributions (KUDs) showing the combined home-range (95% UD; lighter shading) and core (50% UD; darker shading) foraging areas of tracked Tristan skuas during (a) the pre-laying exodus (green), breeding (purple), and non-breeding periods (orange) across the 2017/18 and 2018/19 seasons, overlaid on major oceanic frontal features. Panels (b) and (c) illustrate individual migration routes from the Gough Island region to non-breeding foraging grounds on the South African and Namibian continental shelves, and the corresponding return migrations, respectively. The solid bold red line denotes the movement of one individual (GLS tag BH142) that tested positive for highly pathogenic avian influenza virus (HPAIV) in 2024.

Two of the three dead skuas sampled for HPAIV had been ringed on Gough Island in 2016 and 2017. The individual ringed in 2017 (SAFRING 888666) was a known breeder on the island and had been equipped with a GLS device (BH142) as part of the tracking study (Fig. 4). In contrast, the skua ringed in 2016 (SAFRING 888054) was caught as a non-breeder at the skua club near the helipad.

## Discussion

The arrival of HPAIV H5N1 on Gough Island represents one of the most geographically isolated detections to date, highlighting the interconnectivity of pelagic seabird populations and their role in assisting global transmission in the current HPAIV H5N1 epizootic into some of the world’s most remote (and sensitive) ecosystems. Confirmation of HPAIV in Tristan skuas on Gough prompted suspicions that individuals had become infected by viruses originating from Africa due to the species’ extensive connectivity with southern African coastal habitats outside their breeding season [34, this study]. The GLS data analysed in this study corroborated this migratory pattern, suggesting high temporal and spatial consistency in the species’ use of the Benguela upwelling system as the species’ wintering foraging ground. Multiple outbreaks of HPAIV H5 clade 2.3.4.4b among seabird colonies in southern Africa [18–20, 46] could have provided opportunities for transmission to scavenging species such as skua. However, the phylogenetic analysis of global H5 HA clade 2.3.4.4b sequences places the Gough Island viruses within a cluster of South American strains, distinct from viral strains detected in Africa, and closely related to those found in South Georgia. An expansion of the transmission pathway via South America and the sub-Antarctic is also supported by considering the temporal context of the outbreaks in relation to seasonal and migratory movements within and across species. Based on tracking records for 2018, Tristan skua complete their south-westward migration to return to breeding sites in one to three weeks, arriving near Gough Island between mid-June and beginning of August. While the incubation period for HPAIV can vary considerably depending on species susceptibility and viral genetic characteristics, given the first observation of HPAIV on Gough in mid-September, it is unlikely symptoms would only present 2–4 months after infection (assuming broadly similar migratory timings in 2024). Although we cannot rule out the possibility that the virus arrived first on the African coast before spreading to Gough Island, the timing of infection makes it improbable that the skuas picked up the virus during their winter migration. Therefore, based on both the phylogenetic analysis and outbreak timing, it is more likely that the virus reached Gough via Southern or Atlantic Ocean flyways, potentially during the species’ pre-laying exodus period in August and September, when Tristan skuas disperse into the frontal region south of the island.

While terrestrial aggregations such as breeding colonies represent clear and obvious hotspots for virus transmission, at-sea connectivity may play an equally critical role in transmission to remote locations such as Gough Island. Overlapping foraging grounds and shared marine habitats increase the potential for interspecies contact and facilitating viral spread between geographically distant populations. Although there are currently no known reports of HPAIV transmission occurring at sea, scavenging species are more likely than other seabird species to interact with carcasses, spend extended periods of time sitting on the water, and come into close contact with other foraging individuals [45, 47, 48]. Such behaviours increase opportunities for viral transmission in circumstances where dilution of virions in the environment would otherwise likely preclude infection, thereby supporting the role of skuas and giant petrels as the most likely vectors of AIV across the Southern Ocean [23].

The nutrient-rich boundary waters between the sub-tropical and Antarctic polar fronts are known to support large congregations of pelagic seabirds and serve as important foraging grounds for marine mammals [21, 29, 49]. The high frequency and likelihood of interspecies interactions at these shared foraging grounds, whether through direct or indirect contact, make such zones high-risk areas for the persistence and transmission of infectious diseases. Seabird species breeding on Gough Island use both the Southern and Atlantic Ocean Flyways [21, 32, 50–52], which overlap considerably between 30-60°S, linking Antarctic seabird populations with those on islands in the South Atlantic [21]. In addition, populations breeding on South Georgia have been found to utilise waters near Gough Island including southern elephant seals [53, 54], brown skua [47], and black-browed albatross *Thalassarche melanophris* [31]. Ultimately, however, the timing and location of transmission of HPAIV to the Tristan skuas in this study remain unknown, along with the uncertainty about whether, in fact, skuas introduced the virus to the island or other avian or marine mammal species, either vagrants or those breeding on Gough [50, 55–57].

The detection of HPAIV H5N1 originating from the Antarctic peninsula, in a species regularly foraging in African coastal waters, illustrates a potentially novel eastward trans-oceanic circulation loop, whereby influenza viruses of South American origin could be introduced into wild bird populations of Africa, representing a reverse transmission pathway into the Afro-Eurasian flyway. The Benguela upwelling region, stretching from Cape Agulhas in South Africa north to southwest Angola, is one of the most productive marine ecosystems globally [58] and plays a vital role as a migratory corridor and seasonal feeding area for numerous migratory species breeding in the southern Indian Ocean, Antarctic or sub-Antarctic (e.g. White-capped/Shy albatross (*T. cauta*) [34]; Indian yellow-nosed albatrosses (*T. carteri*) and northern giant petrels (*Macronectes halli*) [59]; black-browed albatrosses [31]) [21]. Therefore, understanding the spatial and temporal dynamics of at-sea dispersal and interactions, especially in areas of high biodiversity and migratory fly- and swim-ways, is essential for anticipating and mitigating future outbreaks.

Initial concerns that HPAIV detection could constitute the onset of a mass outbreak on Gough Island, have not materialized; no further symptomatic birds or mortalities related to HPAIV have been confirmed (aside from a suspicious death of a breeding female Tristan albatross in April and one skua in October 2025). Although mass mortality has occurred in other skua populations as a result of HPAIV, notably in the UK where great skua (*Stercorarius* skua) declined by 73% [4, 45], the apparent failure of the virus to cause a large-scale outbreak on Gough may be due to several factors. Cloacal swabs taken during the initial investigation returned negative results for AIV, suggesting minimal environmental shedding. Additionally, while one carcass was presumed scavenged, the remaining three were buried following sampling, preventing further scavenging. Furthermore, while Gough hosts large populations of seabird species, many of these species nest at low densities (e.g., Tristan albatross) or in burrows, such as petrels and prions [13], reducing opportunities for direct contact, inter-colony mixing and thus, viral spread across the island. A serological study in 2009 demonstrated some seropositivity against influenza in northern rockhopper penguins and Tristan skua on Gough [22, 23]; however, how far pre-existing immunity against AIVs could have influenced transmission dynamics in this outbreak remains unknown.

Detection of outbreaks of HPAIV in wildlife, to infer transmission routes, and understand population impacts, is dependent on reporting of wildlife mortality events and subsequent sampling and testing of carcasses for AIV, with levels of coverage varying substantially across regions. Gaps in surveillance lead to unsampled ancestry in sequence data, limiting the ability to resolve transmission chains from the phylogenetic analysis. However, although surveillance for HPAIV remains limited across many African countries [60], governmental and non-governmental agencies in South Africa and Namibia routinely conduct both active and passive monitoring [18–20, 46, 60, 61]. Phylogenetic analysis of 2023 outbreak sequences in poultry and wild birds shows these strains originated from Europe via West Africa [19]. Viruses sequenced in 2025 are likewise most closely related to the dominant European strain of 2024/2025 [60, L. Roberts pers. comm.], with no South American cluster strains detected in Africa to date. Thus, the strains reported in southern Africa continue to represent a distinct cluster, separate from strains that spread via South America into the Antarctic region and our detection reported here.

Wildlife disease monitoring of populations, particularly in parts of the polar and sub-Antarctic region, currently remain extremely limited. The sheer number of sites and their remote location make any systematic surveillance inherently challenging, and early or subclinical infections in wild or migratory birds or mammals may go unnoticed for extended periods. Avian influenza may have also been present in the Indian Ocean or Pacific sectors of the Southern Ocean before 2024. The absence of detections in these areas, however, may reflect under-sampling rather than true absence, which poses a barrier to early awareness of emerging issues and understanding of patterns of spread.

Although there have been no further mortalities attributed to HPAIV on Gough Island or in the neighbouring Tristan da Cunha Archipelago, the impact of HPAIV in the region could be substantially underestimated. Large parts of this remote group of islands are inaccessible, limiting timely monitoring to detect carcasses or individuals exhibiting clinical signs. While there has been consistent presence of a field team of biologists on Gough year-round since 2008, there is less systematic surveillance at the Tristan da Cunha Archipelago. In February 2024, the island’s fishery observer reported a dead skua floating at sea off Nightingale Island, but no further unusual mortality of skuas (or other seabirds) was observed on Nightingale and Inaccessible islands when visited in March and September 2024, respectively (PGR pers. obs.). Therefore, the absence of confirmed cases should not be taken as evidence of the absence of the disease.

The detection of HPAI on Gough Island is a stark reminder of the vulnerability of remote ecosystems to infectious diseases, and for the need to alleviate already persisting pressures on globally important seabird populations including habitat degradation and disturbance, invasive species, and climate-driven stressors to strengthen their resilience. Proactive and geographically inclusive surveillance strategies and biosecurity measures are vital to understand and mitigate the impacts of this disease, rather than reactive responses after visible outbreaks. Understanding the spatiotemporal dynamics of seabird movements-in particular scavenging seabirds-disease ecology, and host-pathogen interactions is also essential for mitigating the impact of HPAIV and other emerging disease threats, particularly in high-risk and ecologically sensitive areas [62].

## Acknowledgements

All samples for HPAI testing were collected with the permission of the Tristan da Cunha government. We are extremely grateful to Dr Laura Roberts, Western Cape Government and Dr Gretna DeWit, Directorate of Animal Health, South Africa for their invaluable help issuing the South African import permit. A special thank you to Ashley Bennison and Jennifer Forster Davidson, British Antarctic Survey, for their generous and insightful advice and guidance on the Gough HPAIV response plan. We would also like to thank the crews of the *SA Agulhas II* for their kind support and collaboration to receive the samples onboard the vessel. The engagement, shipment to the UK, testing and generation of the viral sequences was funded by the Department for Environment, Food and Rural Affairs (Defra, UK) and the Devolved Administrations of Scotland and Wales, through the following programmes: SV3006, SV3032 and SE2227. This work was also supported by the Biotechnology and Biological Sciences Research Council (BBSRC) and Department for Environment, Food and Rural Affairs (Defra, UK) research initiative ‘FluTrailMap’ [grant number BB/Y007271/1]. Funded by the European Union (EU) under grant agreement (101084171) - (Kappa-Flu). Views and opinions expressed are however those of the author(s) only and do not necessarily reflect those of the EU or REA. Neither the EU nor the granting authority can be held responsible for them. The Science Animal Ethics Committee (SFAEC) of the University of Cape Town approved the research protocols for the deployments of GLS devices on seabirds including skuas (UCT SFAEC 2014\V10\PR), with permission from the Tristan government.

---

**Figure S1:**
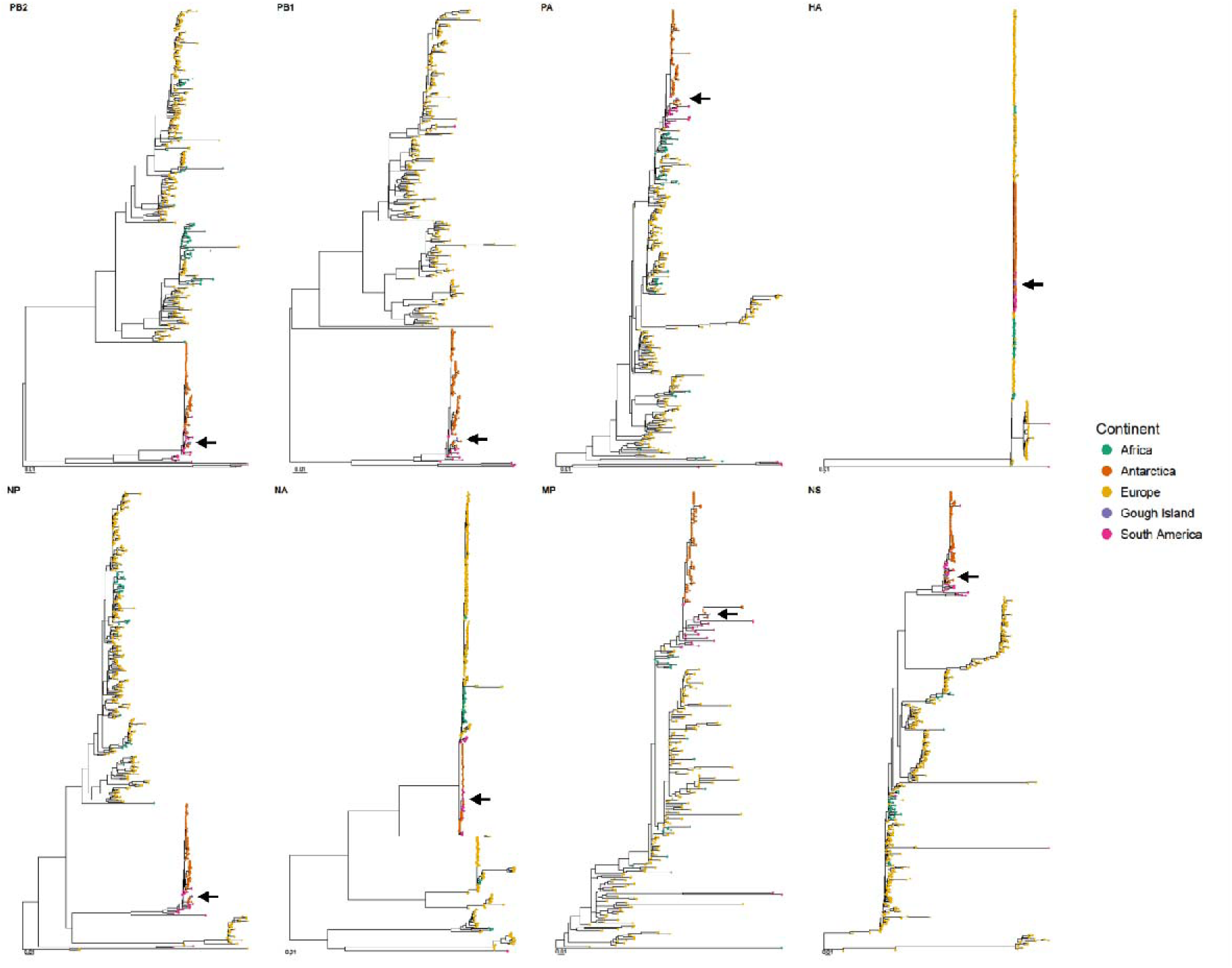
Maximum-likelihood phylogenies of all eight influenza A virus gene segments. Phylogenetic trees were inferred using maximum-likelihood methods for each gene segment show that the Gough Island viruses are closely related to Antarctic H5 viruses rather than South African H5 viruses. Tip shapes represent individual sequences and are coloured according to the continent of origin.

## References

[1] Caliendo V, Lewis NS, Pohlmann A, et al. Transatlantic spread of highly pathogenic avian influenza H5N1 by wild birds from Europe to North America in 2021. Sci Rep. 2022;12(1):11729. doi:10.1038/s41598-022-13447-z

[2] Falchieri M, Reid SM, Ross CS et al. Shift in HPAI infection dynamics causes significant losses in seabird populations across Great Britain. Veterinary Record. 2022;191(7):294–6.

[3] Lane JV, Jeglinski JW, Avery Gomm S, et al. High pathogenicity avian influenza (H5N1) in Northern Gannets (*Morus bassanus*): Global spread, clinical signs and demographic consequences. Ibis. 2024;166(2):633–50. doi:10.1111/ibi.13275

[4] Tremlett CJ, Cleasby IR, Bolton, et al. Declines in UK breeding populations of seabird species of conservation concern following the outbreak of high pathogenicity avian influenza (HPAI) in 2021–2022. Bird Study. 2024;71(4):293–310. doi:10.1080/00063657.2024.2438641

[5] Kuiken T, Vanstreels RE, Banyard A, et al. Emergence, spread, and impact of high pathogenicity avian influenza H5 in wild birds and mammals of South America and Antarctica. Conserv Biol. 2025;e70052. doi:10.1111/cobi.70052

[6] Pardo-Roa C, Nelson MI, Ariyama N, et al. Cross-species and mammal-to-mammal transmission of clade 2.3. 4.4 b highly pathogenic avian influenza A/H5N1 with PB2 adaptations. Nat Commun. 2025;16(1):2232. doi:10.1038/s41467-025-57338-z

[7] Leguia M, Garcia-Glaessner A, Muñoz-Saavedra B, et al. Highly pathogenic avian influenza A (H5N1) in marine mammals and seabirds in Peru. Nat Commun. 2023;14(1):5489. doi:10.1038/s41467-023-41182-0

[8] Gamarra-Toledo V, Plaza PI, Angulo F, et al. Highly pathogenic avian influenza (HPAI) strongly impacts wild birds in Peru. Biol Conserv. 2023;286:110272. doi:10.1016/j.biocon.2023.110272

[9] Banyard AC, Bennison A, Byrne AM, et al. Detection and spread of high pathogenicity avian influenza virus H5N1 in the Antarctic Region. Nat Commun. 2024;15(1):7433. doi:10.1038/s41467-024-51490-8

[10] León F, Ulloa-Contreras C, Pizarro EJ, et al. Skuas mortalities linked to positives HPAIV A/H5 beyond Polar Antarctic Circle. bioRxiv. 2025;1. doi:10.1101/2025.03.02.640960

[11] Clessin A, Briand FX, Tornos J, et al. Mass mortality events in the sub-Antarctic Indian Ocean caused by long-distance circumpolar spread of highly pathogenic avian influenza H5N1 clade 2.3. 4.4 b. 2025. BioRxiv;2025-02. doi:10.1101/2025.02.25.640068

[12] Ryan PG, Gill R, Roberts L, Stephen V. Avian flu reaches Marion Island. African Birdlife. 2025;13(5):14–15.

[13] Caravaggi A, Cuthbert RJ, Ryan PG, et al. The impacts of introduced House Mice on the breeding success of nesting seabirds on Gough Island. Ibis. 2019;161(3):648–61. doi:10.1111/ibi.12664

[14] Oppel S, Clark BL, Risi MM, et al. Cryptic population decrease due to invasive species predation in a long lived seabird supports need for eradication. J Appl Ecol. 2022;59(8):2059–70. doi:10.1111/1365-2664.14218

[15] Jones CW, Risi MM, Osborne AM, et al. Mouse eradication is required to prevent local extinction of an endangered seabird on an oceanic island. Anim Conserv. 2021;24(4):637–45. doi:10.1111/acv.12670

[16] BirdLife International. Rowettia goughensis. The IUCN Red List of Threatened Species 2024: e.T22723149A252044473. 2024 [cited on 07 August 2025]. doi:10.2305/IUCN.UK.2024-2.RLTS.T22723149A252044473.en

[17] Dewar ML, Vanstreels RET, Boulinier T, et al. Biological Risk Assessment of Highly Pathogenic Avian Influenza in the Southern Ocean. Scientific Committee on Antarctic Research. Cambridge UK. SCAR Antarctic Wildlife Health Network. 2023.

[18] Abolnik C, Phiri T, Peyrot B, et al. The molecular epidemiology of clade 2.3.4.4B H5N1 high pathogenicity avian influenza in Southern Africa, 2021–2022. Viruses. 2023;15(6):1383. doi:10.3390/v15061383

[19] Abolnik C, Roberts LC, Strydom C, et al. Outbreaks of H5N1 High Pathogenicity Avian Influenza in South Africa in 2023 Were Caused by Two Distinct Sub-Genotypes of Clade 2.3. 4.4 b Viruses. Viruses. 2024;16(6):896. doi:10.3390/v16060896

[20] Roberts LC, Abolnik C, Waller LJ, et al. Descriptive epidemiology of and response to the high pathogenicity avian influenza (H5N8) epidemic in South African coastal seabirds, 2018. Transbound Emerg Dis. 2023;2023(1):2708458. doi:10.1155/2023/2708458

[21] Morten JM, Carneiro AP, Beal M, et al. Global Marine Flyways Identified for Long Distance Migrating Seabirds From Tracking Data. Glob Ecol Biogeogr. 2025;34(2):e70004. doi:10.1111/geb.70004

[22] Abad, F.X., N. Busquets, A. Sanchez, et al. Serological and virological surveys of the influenza A viruses in Antarctic and sub-Antarctic penguins. Antarct Sci. 2013;25:339–344. doi:10.1017/S0954102012001228

[23] Gittins O, Grau-Roma L, Valle R, et al. Serological and molecular surveys of influenza A viruses in Antarctic and sub-Antarctic wild birds. Antarct Sci. 2020;32(1):15–20. doi:10.1017/s0954102019000464

[24] Ramsar Convention Secretariat. Ramsar Information Sheet (RIS) for Gough Island. Ramsar Sites Information Service; 2025. Site No. 1868. Available from: Ramsar Sites Information Service.

[25] Gamble A, Bazire R, Delord K, et al. Predator and scavenger movements among and within endangered seabird colonies: Opportunities for pathogen spread. J Appl Ecol. 2020;57(2):367–78. doi:10.1111/1365-2664.13531

[26] Sumner MD, Wotherspoon SJ, Hindell MA. Bayesian estimation of animal movement from archival and satellite tags. PLoS One. 2009.13;4(10):e7324. doi:10.1371/journal.pone.0007324

[27] Colchero F, Jones OR, Rebke M. BaSTA: an R package for Bayesian estimation of age specific survival from incomplete mark–recapture/recovery data withcovariates. Methods Ecol Evol. 2012;3(3):466–470. doi: 10.1111/j.2041-210x.2012.00186.x

[28] Calenge C. Home range estimation in R: the adehabitatHR package. Office national de la classe et de la faune sauvage: Saint Benoist, Auffargis, France. 2011.

[29] Weimerskirch H. Are seabirds foraging for unpredictable resources? Deep Sea Res II: Top Stud Oceanogr. 2007;1;54(3-4):211–23. 10.1016/j.dsr2.2006.11.013

[30] Orben RA, Kokubun N, Fleishman AB, et al. Persistent annual migration patterns of a specialist seabird. Marine Ecology Progress Series. 2018 Apr 12;593:231–45. 10.3354/meps12459

[31] Phillips RA, Silk JR, Croxall JP, et al. Summer distribution and migration of nonbreeding albatrosses: individual consistencies and implications for conservation. Ecology. 2005;86(9):2386–96. doi:10.1890/04-1885

[32] Green CP, Green DB, Ratcliffe N, et al. Potential for redistribution of post moult habitat for *Eudyptes* penguins in the Southern Ocean under future climate conditions. Glob Chang Biol. 2023;29(3):648–67. doi:10.1111/gcb.16500

[33] Lewin PJ, Wynn J, Arcos JM, et al. Climate change drives migratory range shift via individual plasticity in shearwaters. Proc Natl Academ Sci, 2024;121(6), e2312438121.

[34] Carneiro APB, Dias MP, Clark BL, et al. The BirdLife Seabird Tracking Database: 20 years of collaboration for marine conservation. Biol Conserv. 2024;299:110813. doi:10.1016/j.biocon.2024.110813

[35] James J, Seekings AH, Skinner P, et al. Rapid and sensitive detection of high pathogenicity Eurasian clade 2.3. 4.4 b avian influenza viruses in wild birds and poultry. J Virol Methods. 2022;301:114454. doi: 10.1016/j.jviromet.2022.114454

[36] Nagy A, Černíková L, Kunteová K, et al. A universal RT-qPCR assay for “One Health” detection of influenza A viruses. PloS one. 2021;16(1):e0244669. doi:10.1371/journal.pone.0244669

[37] Payungporn S, Chutinimitkul S, Chaisingh A, et al. Single step multiplex real-time RT-PCR for H5N1 influenza A virus detection. J Virol Methods. 2006;131(2):143–7. doi:10.1016/j.jviromet.2005.08.004

[38] Sutton DA, Allen DP, Fuller CM, et al. Development of an avian avulavirus 1 (AAvV-1) L-gene real-time RT-PCR assay using minor groove binding probes for application as a routine diagnostic tool. J Virol Methods. 2019;265:9–14. doi:10.1016/j.jviromet.2018.12.001

[39] Slomka MJ, Reid SM, Byrne AM, et al. Efficient and informative laboratory testing for rapid confirmation of H5N1 (Clade 2.3. 4.4) high-pathogenicity avian influenza outbreaks in the United Kingdom. Viruses. 2023;15(6):1344. doi:10.3390/v15061344

[40] Markin A, Wagle S, Grover S, et al. PARNAS: objectively selecting the most representative taxa on a phylogeny. Syst Biol. 2023;72(5):1052–63. doi:10.1093/sysbio/syad028

[41] Katoh K, Standley DM. MAFFT multiple sequence alignment software version 7: improvements in performance and usability. Mol Biol Evol. 2013;30(4): 772–780. doi:10.1093/molbev/mst010

[42] Larsson A. AliView: a fast and lightweight alignment viewer and editor for large datasets. Bioinformatics 30.22. 2014:3276–3278. doi:10.1093/bioinformatics/btu531

[43] Minh BQ, Schmidt HA, Chernomor O, et al. IQ-TREE 2: new models and efficient methods for phylogenetic inference in the genomic era. Mol Biol Evol 2020;37(5):1530–1534. doi:10.1093/molbev/msaa015

[44] Sagulenko P, Puller V, Neher RA. TreeTime: Maximum-likelihood phylodynamic analysis. Virus Evol. 2018;4(1):vex042. doi:10.1093/ve/vex042

[45] Banyard AC, Lean FZ, Robinson C, et al. Detection of highly pathogenic avian influenza virus H5N1 clade 2.3. 4.4 b in great skuas: a species of conservation concern in Great Britain. Viruses. 2022;14(2):212. doi:10.3390/v14020212

[46] World Organisation for Animal Health (WOAH), 2025. WAHIS Event 5034 Influenza A viruses of high pathogenicity (Inf. with) (non-poultry including wild birds) (2017-). https://wahis.woah.org/#/in-event/5034/dashboard

[47] Carneiro AP, Manica A, Clay TA, et al. Consistency in migration strategies and habitat preferences of brown skuas over two winters, a decade apart. Mar Ecol Prog Ser. 2016;553:267–81. doi:10.3354/meps11781

[48] Sinclair JC. Subantarctic Skua *Catharacta antarctica* predation techniques on land and at sea. Mar Ornithol. 1980;8:3–6. doi:10.5038/2074-1235.8.1.27

[49] Bost CA, Cotté C, Bailleul F, et al. The importance of oceanographic fronts to marine birds and mammals of the southern oceans. J Mar Syst. 2009;78(3):363–376. doi:10.1016/j.jmarsys.2008.11.022

[50] Dias MP, Oppel S, Bond AL, et al. Using globally threatened pelagic birds to identify priority sites for marine conservation in the South Atlantic Ocean. Biol Conserv. 2017;211:76–84. doi:10.1016/j.biocon.2017.05.009

[51] Schoombie S, Dilley BJ, Davies D, et al. The foraging range of Great Shearwaters (*Ardenna gravis*) breeding on Gough Island. Polar Biol. 2018;41(12):2451–8. doi:10.1007/s00300-018-2381-7

[52] Jones CW, Phillips RA, Grecian WJ, et al. Ecological segregation of two superabundant, morphologically similar, sister seabird taxa breeding in sympatry. Mar Biol. 2020;167(4):45. 10.1007/s00227-020-3645-7

[53] Reisinger RR, Bester MN. Long distance breeding dispersal of a southern elephant seal. Polar Biol. 2010;33(9):1289–91. doi:10.1007/s00300-010-0830-z

[54] Hindell MA, McMahon CR, Bester MN, et al. Circumpolar habitat use in the southern elephant seal: implications for foraging success and population trajectories. Ecosphere. 2016;7(5):e01213. doi:10.1002/ecs2.1213

[55] Bester MN, Wege M, De Bryn, et al. Ranging and fiving behaviour of Subantarctic fur seals from the Tristan da Cunha Islands, South Atlantic. Tristan da Cunha Islands Seal Research Report (2009 – 2019). 2019

[56] Ryan PG, Dilley BJ, Risi MM, et al. Three new seabird species recorded at Tristan da Cunha archipelago. Seabird. 2019;32:122–5. doi:10.61350/sbj.32.122

[57] Bester MN, Somers MJ. Monitoring of vagrant seals on mid-oceanic islands of the South Atlantic Ocean. Polar Biol. 2025;48(3):79. doi:10.1007/s00300-025-03397-3

[58] Hutchings L, Van der Lingen CD, Shannon LJ, et al. The Benguela Current: An ecosystem of four components. Prog in Oceanogr. 2009;83(1-4):15–32. doi:10.1016/j.pocean.2009.07.046

[59] Reisinger RR, Raymond B, Hindell MA, et al. Habitat modelling of tracking data from multiple marine predators identifies important areas in the Southern Indian Ocean. Divers Distrib. 2018;24(4):535–50. doi:10.1111/ddi.12702

[60] Abolnik C. Avian influenza situation report—Africa. Canadian Journal of Microbiology. 2025;26;71:1–4. 10.1139/cjm-2025-0199

[61] Molini U, Yabe J, Meki IK, et al. Highly pathogenic avian influenza H5N1 virus outbreak among Cape cormorants (Phalacrocorax capensis) in Namibia, 2022. Emerg Microbes Infection. 2023;12(1):2167610. 10.1080/22221751.2023.2167610

[62] Sacristán C, Ewbank AC, Ibáñez Porras P, et al. Novel epidemiologic features of high pathogenicity avian influenza virus A H5N1 2.3. 3.4 b panzootic: a review. Transbound Emerg Dis. 2024;2024(1):5322378. doi:10.1155/2024/5322378

